# A discovery pipeline for identification and *in vivo* validation of drugs that alter T cell/ dendritic cell interaction

**DOI:** 10.1101/2022.04.11.487526

**Authors:** Hannah E. Scales, Chantal Keijzer, Emma Evertsson, Paul Garside, Stephen Delaney, James M. Brewer

## Abstract

The interaction between T cells and dendritic cells (DC) is central to determining whether or not an adaptive immune response develops. Autoimmune diseases like rheumatoid arthritis result from inappropriate activation of self-antigen specific T cells, therefore targeting the interaction between T cells and DCs is of considerable therapeutic interest. Here, we describe the development and validation of a drug development platform that includes a high-throughput image-based screening assay and a complementary *in vivo* model. The *in vitro* assay directly measures the ability of compounds to influence the interaction of T cells with DC, both negatively and positively. While the *in vivo* model allows the validation of hit molecules directly in the lymph nodes of intact animals. Using this pipeline, we identify a new function for the protein kinase inhibitor AS703569 by inhibiting T cell/ DC interactions *in vitro and in vivo*.

## INTRODUCTION

The immune system is composed of a complex group of organs, cells and molecules which collectively function to recognise and remove pathogens or abnormal host cells, while simultaneously leaving harmless environmental antigens and healthy host tissue untouched. The decision to activate in response to pathogens or induce tolerance to self-antigens is made through the antigen specific interactions of T cells with dendritic cells (DC). The nature of these interactions has been shown to be directly associated with immunological outcomes; with long stable interactions driving immune activation, while shorter interactions favour tolerance (1–3). T cell/DC (T/DC) interactions involve the formation of a highly organised multi-molecular structure at the interface of the T cell and the DC, known collectively as the immunological synapse (IS)(4, 5). In addition to cognate interactions between the T Cell Receptor (TCR) and peptide Major Histocompatibility Complex (MHC), the formation of the IS requires the interaction of multiple surface molecules and intracellular signalling pathways in both cell types (6). For example, both the T cell and the DC undergo extensive cytoskeletal reorganisation (7, 8); activation of the TCR leads to intracellular signalling via Lymphocyte-specific protein tyrosine kinase (LCK) and multiple intracellular kinases such as C-Terminal Src Kinase (CSK) and the Mitogen-activated protein (MAP) Kinases (9, 10); while the interaction between the costimulatory molecule CD40 expressed on the DC cell surface with CD40 ligand (CD40L) expressed on the T cell surface leads to activation of both the DC and the T cell (11–13).

Targeting the IS for therapeutic intervention has proven difficult due to the complexity, redundancy and pleiotropic nature of the pathways and molecules involved. Exceptions to this are Abatacept, a Cytotoxic T Lymphocyte Antigen-4-immunoglobulin (CTLA-4-Ig) fusion protein, and the checkpoint inhibitors anti-CTLA-4 and anti-Programmed Cell Death Protein 1 (PD1). Abatacept prevents engagement of the costimulatory molecule CD28 on the T cell surface with its activating ligands expressed by DC there by inhibiting T cell activation (14) and is used clinically to treat Rheumatoid arthritis (RA) patients (15–17). Checkpoint inhibitors conversely prevent the inhibition of T cell and other immune cell activation by cancer cells and have been approved for the use in a number of otherwise difficult to treat malignant diseases (18).

Currently, *in vitro* assays investigating T/DC interactions tend to rely largely on either reporters of specific molecules downstream of the TCR; the up or down regulation of T cell activation markers. As a result, these assays are biased, requiring prior knowledge of the target pathway or may show effects that are not functionally related to T cell activation.

In order to address this, we have developed a functional, live cell-based screening assay that utilises high-content, high-throughput microscopy to directly assess T/DC interaction *in vitro* in an unbiased fashion. This *in vitro* assay is complemented by an *in vivo* model where we employ multiphoton laser scanning microscopy (MPLSM) to directly measure the effect of a compound on the T/DC interaction *in situ* in an intact lymph node (LN). Using this pipeline, we have screened 151 members of a kinase inhibitor library. In addition to validating the assay through identification of kinases involved in T cell activation, we have also identified a new function for an existing kinase inhibitor (AS703569) through inhibition of T/DC interaction *in vitro* and *in vivo*. These studies demonstrate the utility of the image-based assay as an efficient *in vitro* screening platform with direct *in vivo* translation.

## RESULTS

### An image based high-throughput screening assay assessing T/DC interactions

In order to optimise the culture conditions to identify the interaction of T cells with DCs we co-cultured fluorescently labelled bone marrow derived DCs (BMDC) and purified CD4^+^ OTII T cells in 384 well plates. These cells were co-cultured at a DC:T ratio of 1:1 or 1:2 without antigen, with increasing concentrations of Ovalbumin peptide (pOVA); with whole OVA protein or with the non-specific T cell mitogen Concanavalin A (ConA). Representative images show increased clustering, which is indicative of T/DC interaction, in the presence of pOVA and ConA (Figure 1A). Analysis of the images obtained demonstrate that the percentage of the area of CFSE (T cell) staining that overlaps with CMTPX signal (DC) increases with the addition of OVA peptide, whole OVA protein and in the presence of ConA compared with media alone (Figure 1B and C); indicating that this is a valid measure of T/DC interaction. Neither increasing the ratio of T cells to DCs or the overall number of cells per well altered the degree of interaction observed (Figure 1B), however increasing the concentration of pOVA from 0.1 to 1 μg/mL did significantly increase interaction, however increasing the pOVA concentration further did not (Figure 1C).

**Figure 1.**
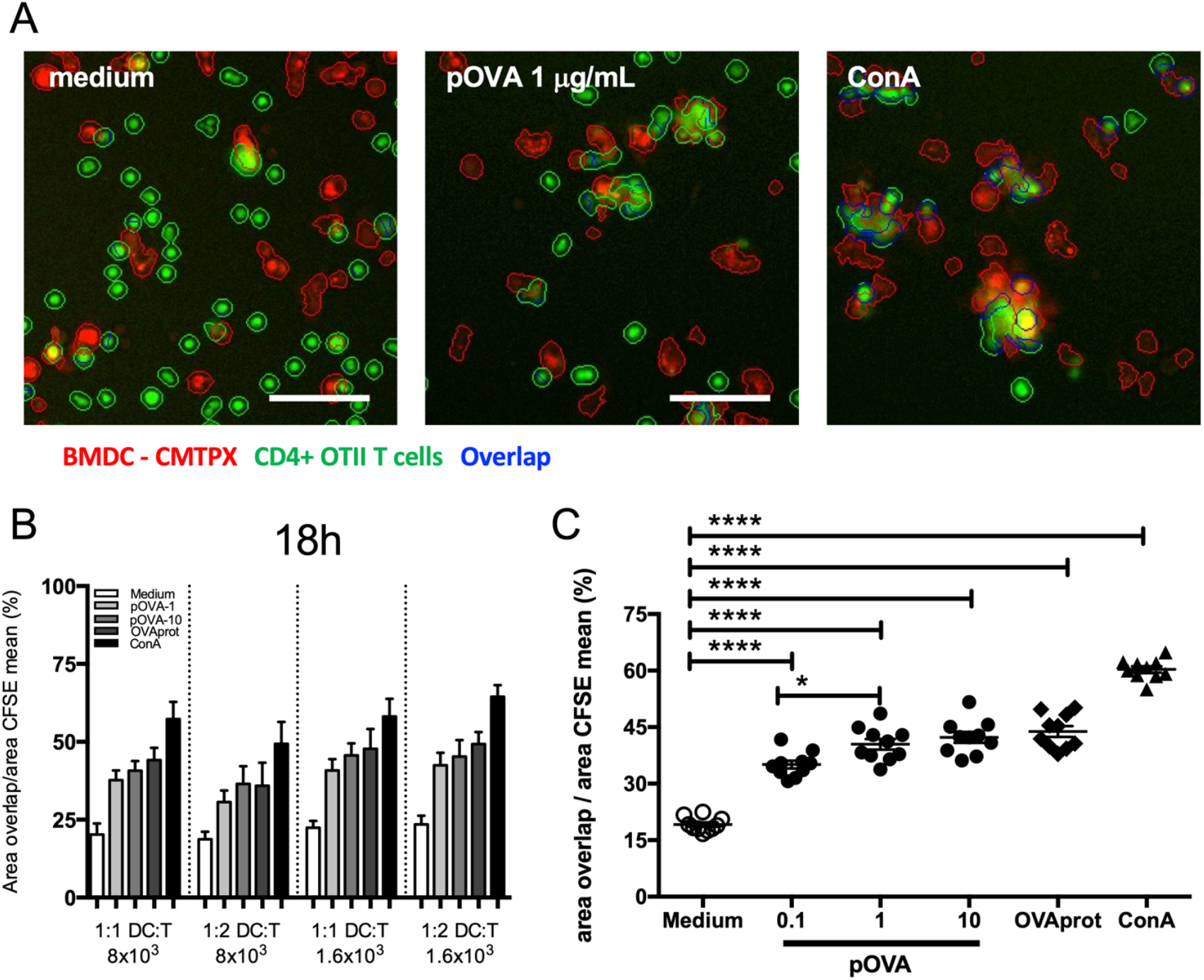
Optimised live cell-based screening assay that utilises high-content, high-throughput microscopy to directly assess T/DC interaction in vitro in an unbiased fashion. 8000 or 16000 purified OTII CD4+ T cells (CFSE, green) were co-cultured in clear bottom 384 well plates with fluorescently labelled bone marrow derived DC (CMTPX, red) at DC:T ratios of 1:1 and 1:2. Cell-cell interaction was stimulated with the T cells cognate antigen either as the specific peptide (pOVA) at 0.1, 1 and 10 μg/mL or whole OVA protein at 100 μg/mL. ConA was used at 2.5 μg/mL as a non-specific stimulator of interaction. The cultures were incubated for 18 hours and the wells imaged on an INCell Analyzer 2000. A shows representative images from unstimulated (medium); and cultures stimulated with either pOVA at 1 μg/mL or ConA at 2.5 μg/mL at a 1:1 DC:T ratios. Scale bar = 50 μm. B and C show quantification of cellular overlap calculated as a percentage of the area of CFSE^+^ T cells that overlap with the CMTPX labelled DCs. Graphs show mean ± SEM and statistical significance was determined using a One-way ANOVA with Tukey multiple comparisons test. *P<0.05, **** P<0.0001.

### T/DC interaction in both antigen or mitogen stimulated cultures can be positively or negatively modulated

In order to determine the stimulation conditions that are best for identifying molecules that alter T/DC interactions we employed LPS as a molecule known to enhance T/DC interactions *in vivo* and an MHCII specific antibody Y-3P, which blocks the interaction of T cells with APCs *in vivo* (1, 19). Labelled T cells and DC were cultured with pOVA or ConA at a range of concentrations (0.01 to 2.5 μg/mL) with or without LPS at 100 ng/mL or Y-3P at 1 μg/mL. Both pOVA and ConA stimulated T/DC interaction in a dose dependant fashion, LPS stimulated an increase in the pOVA induced interaction at almost all the pOVA concentrations tested (Figure 2A). ConA at 2.5 μg/mL appears to maximise interaction, however LPS increased interaction at the ConA concentrations from 0.1 to 1 μg/mL (Figure 2B). As expected, Y-3P treatment completely ablated antigen driven (pOVA) T/DC interaction (Figure 2C); Y3P treatment was also able to block ConA driven interaction although not to the same extent (Figure 2D). By utilising a known adjuvant (LPS) and an antibody known to block T/DC interaction these data demonstrate the potential for this assay to identify compounds that either inhibit (immunemodulators) or enhance (adjuvants or checkpoint inhibitors)T/DC interaction.

**Figure 2.**
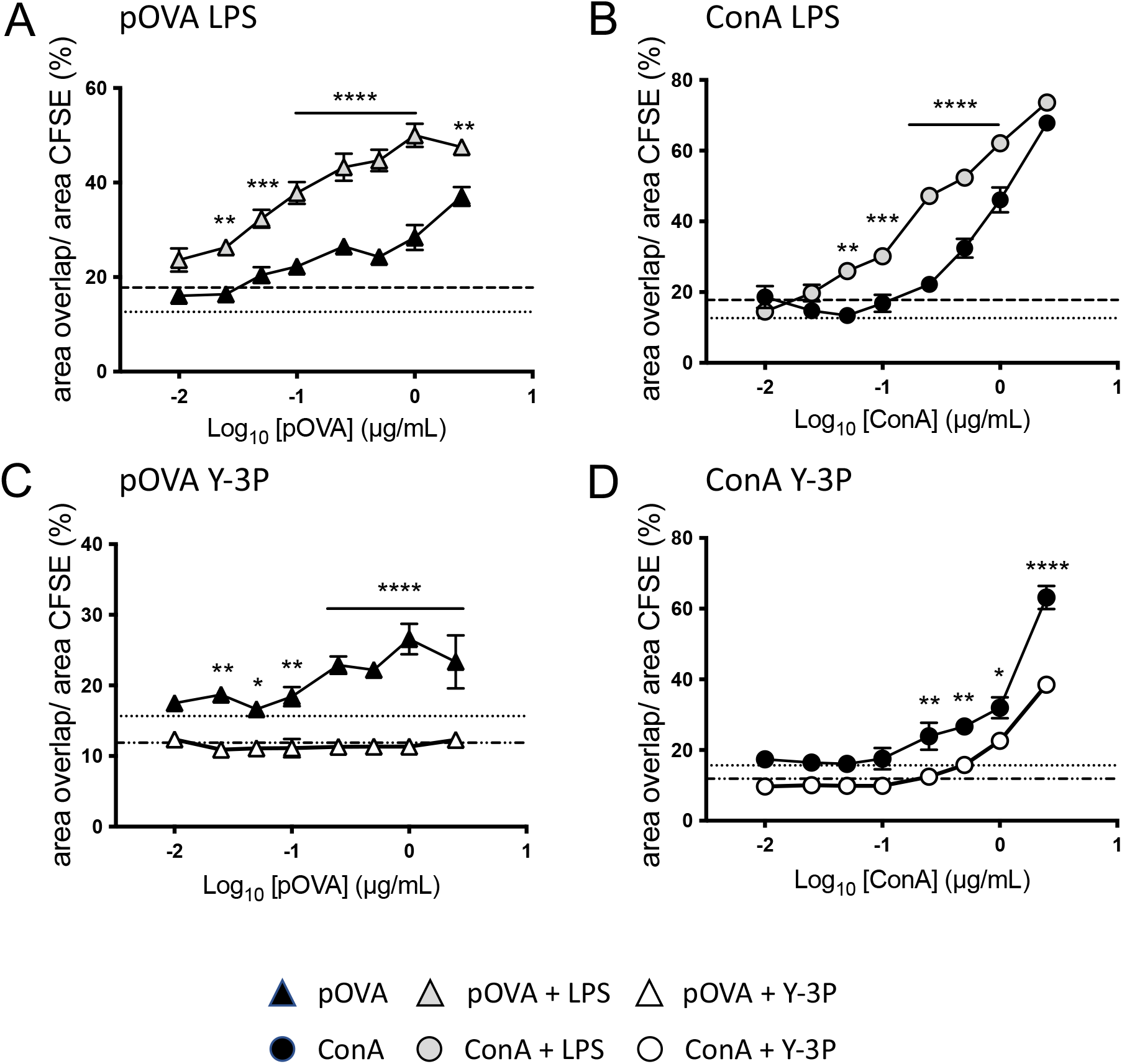
Antigen-specific and non-specific T/DC interactions can be enhanced with an adjuvant or inhibited with an MHCII blocking antibody. Fluorescently labelled BMDC (CMTPX, red) and purified OTII CD4^+^ T cells (CFSE, green) were co-cultured in clear bottom 384 well plates at a ratio of 1:1. Interaction was stimulated with pOVA or ConA at a range of concentrations (0.1 to 2.5 μg/mL). Cultures were treated with LPS as a prototypical adjuvant at 100 ng/mL or with the MHCII blocking antibody Y-3P at 1 μg/mL. Graphs show the mean percentage overlap against the log concentration of the stimulant. (A) pOVA and (B) ConA stimulated cultures treated with LPS. (C) pOVA and (D) ConA stimulated cultures treated with Y-3P. Graphs show mean ± SEM and statistical significance was determined using a two-way ANOVA with Sidak’s multiple comparisons test. *P<0.05, **P<0.01, *** P<0.001 **** P<0.0001.

### 2.2 Compound library screening identifies compounds with the potential to inhibit T cell –DC interaction in vitro

In order to further validate this assay, we chose 39 compounds with known biochemical inhibitory activity against 43 different kinases listed in Figure 3A (kinase inhibitors have overlapping activity) but unknown activity at the level of T/DC interaction. These compounds and a DMSO vehicle control were plated onto 384 well plates at final well concentration of 10 μM (0.1% DMSO). T cells and DCs were added at 1:1 ratio (8000 of each) and stimulated with either media alone; pOVA at 1 ug/mL or ConA at 2.5 μg/mL. Following overnight incubation, 14 compounds were identified that reduced the T/DC interaction stimulated by both pOVA and ConA (Figure 3B). Inhibitory compounds 8, 9, 10, 17 and 35 and the non-inhibitory compound number 3 were selected for IC50 determination (Figure 4A and B). Compounds were further assessed by flow cytometry for their effect on the viability of splenocytes cultured with the compounds overnight. Compound 35 was found to have the lowest IC50 and further analysis demonstrated that it did not affect cell viability at this concentration (Figure 4C). Compound 35 was identified as AS703569 and a literature search showed that this compound had been tested as an anti-proliferative agent in tumour models *in vivo* and was well tolerated by mice (20), in combination with the favourable IC50 and viability data we selected this compound for further study.

**Figure 3.**
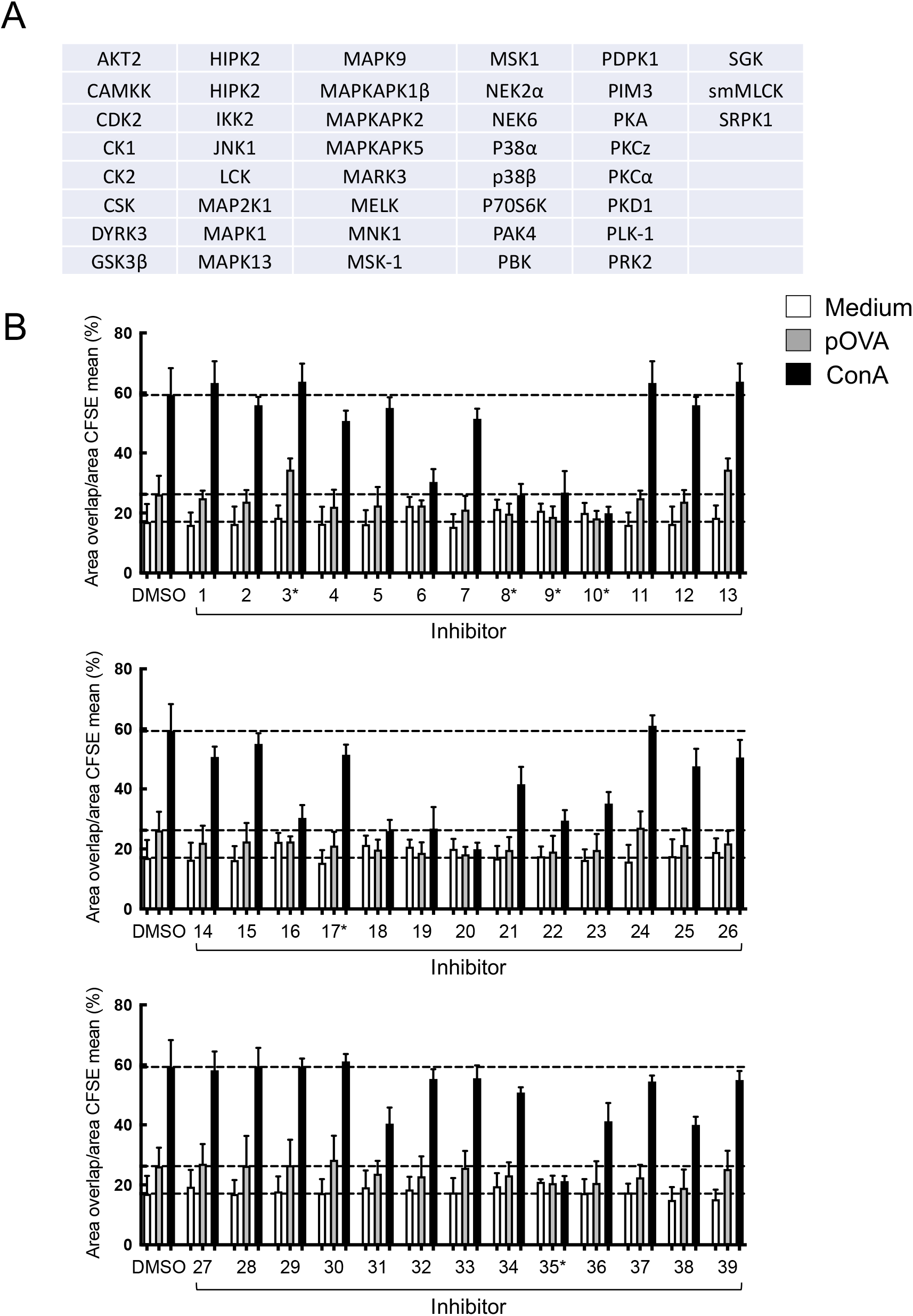
Screening a 39 compound library of kinase inhibitors identifies 14 compounds with apparent activity. (A) Table showing biochemical activity of kinases inhibitor library selected for study. (B) Fluorescently labelled BMDC (CMTPX, red) and purified OTII CD4^+^ T cells (CFSE, green) were cultured in 384 well plates pre-printed with a kinase inhibitor library solubilised in DMSO at 10 μM. Interactions were stimulated with pOVA at 1 μg/mL or ConA at 2.5 μg/mL. Wells were imaged at approximately 18 hours and the degree of interaction was determined from the images. Data is presented as mean overlap ± SEM. Dashed lines represent the mean overlap for ConA, pOVA and unstimulated (medium) treated wells. * Denotes the inhibitors selected for IC50 determination.

**Figure 4.**
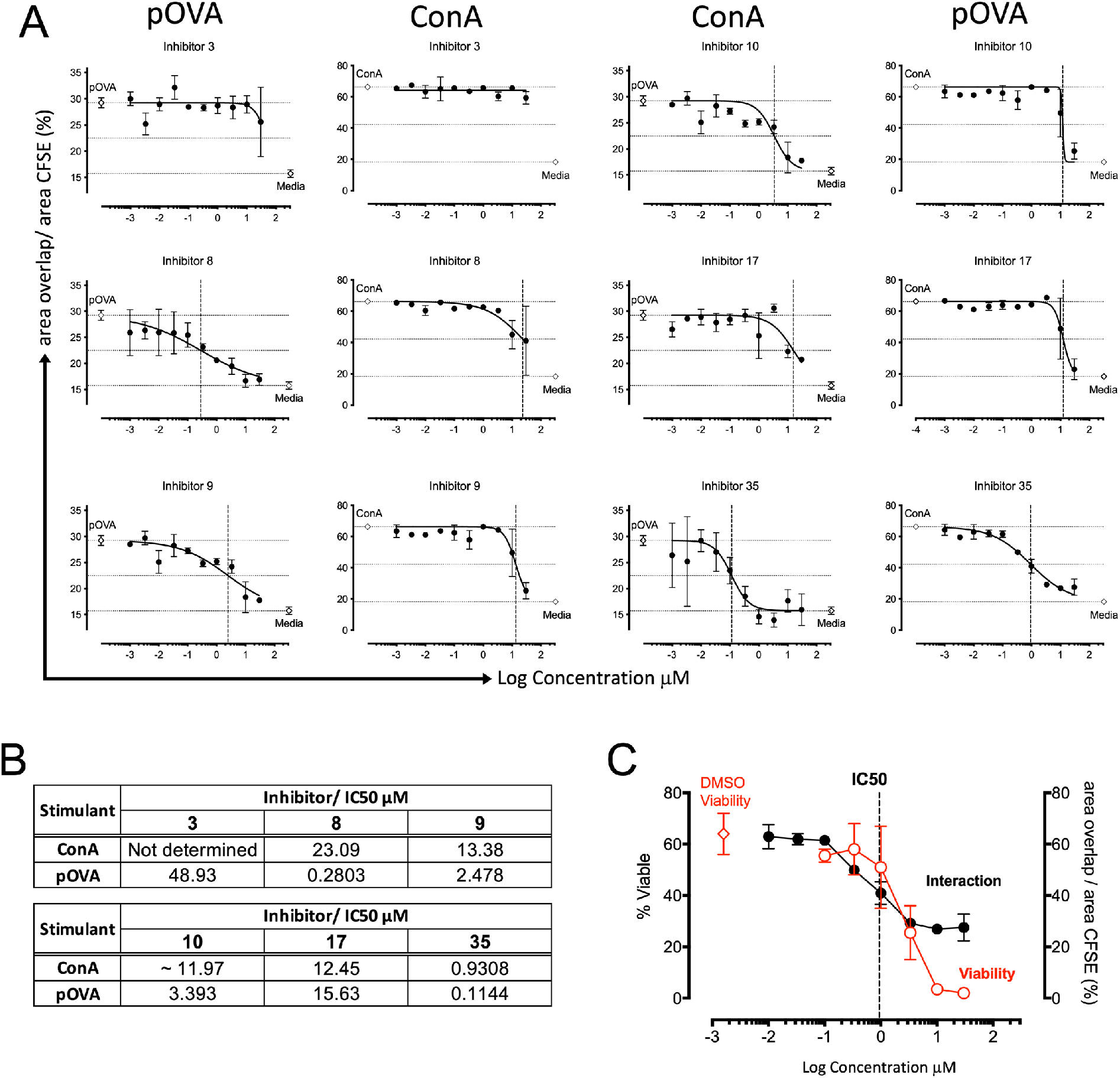
IC50 calculation and viability assessment identified the kinase inhibitor AS703569. (A) A dose response assay (0.0001 to 30 μM) was performed using six compounds selected from the initial screen in the pOVA and ConA stimulated T/D interaction assay and the IC50 for each compound/stimulation calculated. The IC50 is marked as a dashed line on each graph and in Table B. (C) The effect of compound 35 on cell viability was assessed by flow cytometry on *in vitro* splenocyte cultures.

### 2.3 AS703569 inhibits T cell activation in vitro and T/DC interaction in vivo

T/DC co-cultures treated with AS703569 at 0.3 μM show the inhibition of T cell activation markers (reduced CD25 and CD69 and increased CD62L compared to the vehicle control), in response to both pOVA and ConA stimulation at 24 hours (Figure 5A). AS703569 also inhibited T cell proliferation as assessed by CFSE dilution and flow cytometry at 72 hours (Figure 5B).In order to determine whether the *in vitro* screening assay can be used to select compounds that also show activity *in vivo* we tested AS703569 in an *in vivo* model that allows us to directly measure T/DC interaction *in situ* in an intact LN. We adoptively transferred purified CD4^+^ T cells from OTII T cell transgenic mice expressing the fluorescent protein DSRed under the control of the hCD2 promotor into mice expressing the YFP under the control of the CD11c promoter. Mice were treated locally in the foot pad with 5 μg AS703569 and immunised with 100 μg OVA protein and 10 μg LPS. The draining popliteal LN was surgically exposed and imaged *in situ* by multiphoton microscopy. T/DC interactions were imaged, and the duration of interaction was assessed. Immunisation with OVA/LPS increased the duration of interaction and this was abolished by treatment with AS703569 (Figure 5C, 5D and supplementary movies 1 and 2). Interestingly, we observed that while AS703569 significantly reduced the interaction duration of T cells with DCs following immunisation with OVA/LPS, we could also show this effect was relatively short lived. Subsequent movies in the same animal demonstrated that interactions returned to levels similar to vehicle control within (approximately 2 hours) (Supplementary Figure 1). This finding is consistent with AS703569 being a competitive kinase inhibitor and its relatively rapid displacement in the active site by ATP. It also emphasises the utility of the *in vivo* approach to rapidly assess pharmacodynamics prior to further *in vivo* investigations of drug effects.

**Figure 5.**
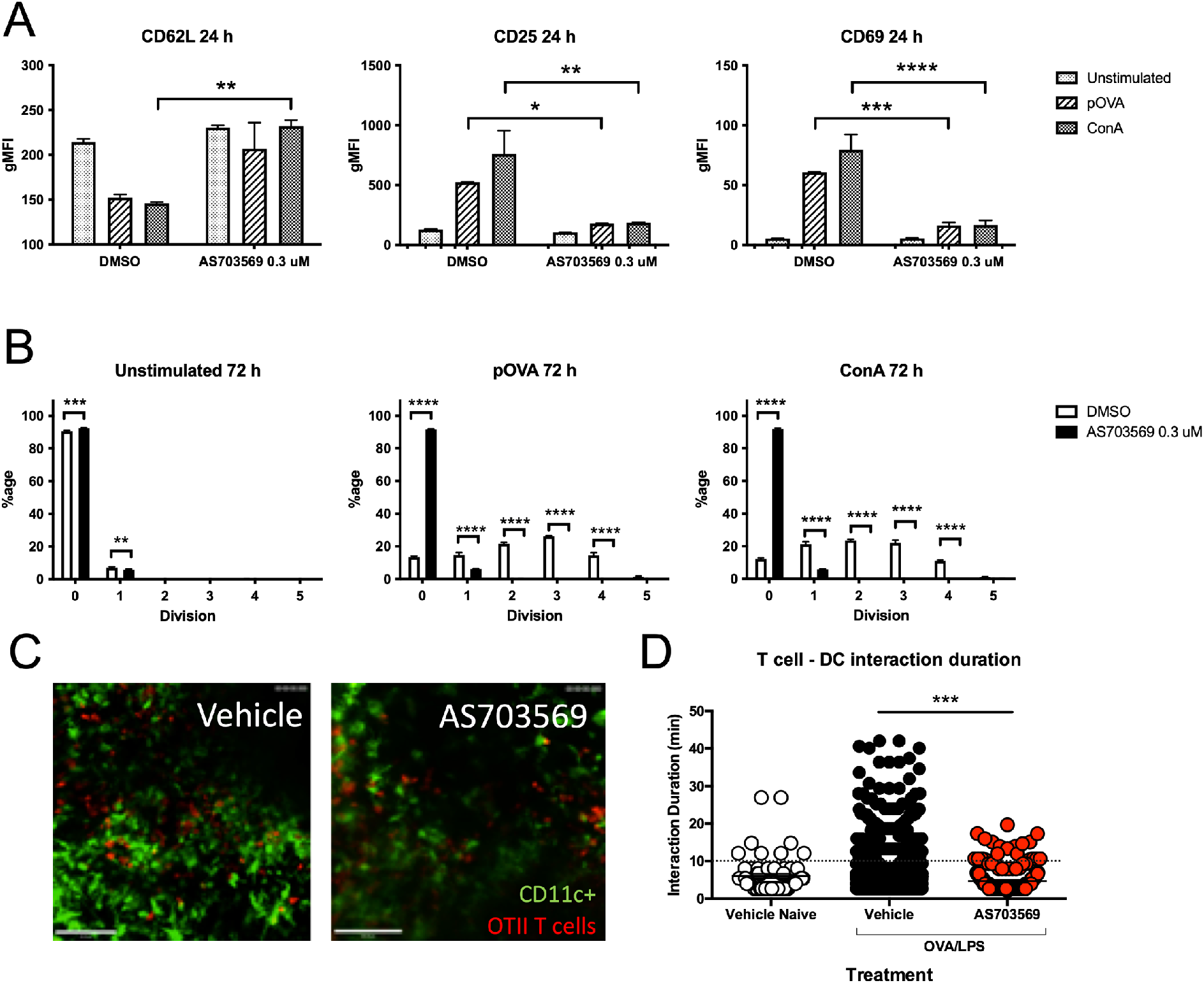
AS703569 inhibits T cell proliferation and activation in vitro and T/DC interaction in vivo. CFSE labelled OTII CD4^+^ T cells and DCs were cultured *in vitro* with and without AS703569 at 0.3 μM. Cultures were stimulated with either pOVA or ConA and the expression of the T cell activation markers CD62L, CD25 and CD69 were measured by flow cytometry at 24 hours (A) T cell proliferation was assessed by measuring CFSE dilution at 72 hours (B). The effect of *in vivo* treatment on T/DC interaction was measured in the popliteal LN of CD11c-YFP mice following the adoptive transfer of DSRed OTII CD4^+^ T cells and subsequent immunisation with OVA/LPS and treatment with 5 μg AS703579 or vehicle control in the footpad. Unimmunised, vehicle treated mice were used as a control. (C) representative stills from movies of vehicle and AS703569 treated mice, DCs are shown in green and T cells in red, scale bar = 100 μm. (D) interaction duration between T cells and DCs in the three treatment groups (naïve n=1 mouse, Vehicle and AS703569 treated n= 3 mice/group). Data from only the first movie was used to generate the graph, each point represents an individual cell. Data is presented as mean ± SEM and statistical significance between the immunised AS703569 and vehicle treated mice was determined using an unpaired T test. *** P<0.001.

This data demonstrates the *in vivo* translate-ability of the *in vitro* screening assay.

## DISSCUSSION

Here, we describe an unbiased, *in vitro* screening assay for the identification of molecules that either enhance or inhibit T/DC interactions. We applied this assay to screen a library of protein kinase inhibitors with unknown effects on T/DC interaction and identified a molecule (AS703569) that inhibits T/DC interaction *in vitro* and most importantly *in vivo*.

By measuring the effect of test compounds on T/DC interaction directly rather than using surrogate markers such as T cell proliferation or activation markers this platforms offers substantial refinement to the process of screening molecules that may affect T cell activation. By focussing on the interaction of cells, the assay is unbiased as prior knowledge of the molecular target is not required. This also negates the requirement for specific molecular reporters. In combination with the *in vivo* T/DC analysis, the number of animals required to establish the function of compounds is reduced.

AS703569 (also known as R763) is a small molecule kinase inhibitor that has previously been identified as a competitive aurora kinase B inhibitor (20). The mammalian Aurora kinase (Aur) family, comprised of three molecules A, B and C, are serine/threonine kinases that regulate processes during cell division such as centrosome formation, spindle dynamics and cytokinesis. An inhibitor of AurB activity may therefore be expected to have an impact on T cell proliferation independently of effects on the IS. However, as this study assessed directly effects on T/DC interaction both *in vitro* and *in vivo* it suggests that AS703569 is affecting pathways responsible for the development of a stable synapse rather than pathways simply inhibiting cell division. In addition, a role for AurA in early TCR signalling, IS formation and T cell activation has been demonstrated using the selective AurA inhibitor MLN8237(21). Taken together these studies suggest that Aurora kinases have the potential to play more complex roles in T cell activity than simply regulating cell proliferation. In light of their role in regulating the activity of components of the cell cytoskeleton and the importance of the cytoskeleton in the formation of a stable IS this is perhaps not a surprising finding. However, in biochemical assays AS703569 has be shown to have activity against a large number of kinase signalling molecules including molecules that are known to function downstream of the TCR signalling pathway such as LCK or on the antigen presenting cell (APC) side of the synapse down stream of CD40 signalling such as JNK (data not shown, AstraZeneca). Therefore, the precise mechanism(s) by which this molecule is preventing the formation of stable T/DC interactions is not clear. AS703569 has be tested in phase I trials for malignant disease where it has been well tolerated and some patients have shown signs of clinical improvement. The generalised antiproliferative effects of inhibiting aurora kinases may make these a difficult target in autoimmune disease. However, applying pharmaceutical chemistry to AS703569 to generate novel compounds and utilising the High Throughput Screening assay may allow separation of these effects and generate further lead compounds.

ConA, a plant derived lectin has been used for many years to stimulate T cell activation, and is known to function by cross-linking surface molecules between T cells and APCs (22). Y-3P is an antibody targeting the MHCII on the DC and has been shown to block the recognition and binding of cognate antigen through the TCR (1, 19). The ability of the antibody Y-3P to significantly reduce ConA induced T/DC interaction in the *in vitro* assay suggests that ConA is cross linking the TCR with MHCII as has been seen previously with CD8+ T cells and MHCI (22). Furthermore, the small molecule kinase inhibitors that prevented antigen (pOVA) driven interactions also tended to have some activity blocking T/DC interactions stimulated by ConA. Conversely, the addition of LPS, which stimulates DCs to increase expression of MHCII and costimulatory molecules therefore demonstrates adjuvant like properties, enhanced T/DC interactions in this assay in response to both specific antigen and ConA. Therefore, this assay has the potential to identify molecules with either immune inhibiting or enhancing properties. Additionally, the non-specific nature of ConA stimulation lends itself to the adaptation of this assay to other species where TCR transgenic cells are not available.

In conclusion, the identification of AS703569 through our assay demonstrates the utility of an unbiased assay within this *in vitro* to *in vivo* platform to identify potential drug compounds with potentially novel modes of action to manipulate T/DC interaction for therapeutic intervention. While small molecule inhibitors have been used to elucidate the function of different actors in biological processes the dynamic interaction between inhibitors, particularly competitive inhibitors and their targets can make their efficacy difficult to assess using *in vivo* disease models. However utilising intravital LN imaging we were able to determine that AS703569 was functional *in vivo* but that the activity was comparatively short lived. The studies presented here highlight the value not only of an *in vitro* high throughput imaging screen to identify immunotherapeutic compounds, but the importance of pairing this approach with a complementary *in vivo* screen to identify the *in vivo* functionality of identified compounds.

## MATERIALS AND METHODS

### Mice

Female C57BL/6J wild-type mice (6-8 weeks) were purchased from Harlan Laboratories, UK. Ovalbumin (OVA) peptide (323-339)-specific TCR transgenic (Tg) (23) (OTII) x CD45.1 mice on a C57BL/6 background were used as a source of OVA specific T cells for the *in vitro* experiments. For the *in vivo* imaging experiments CD11cYFP mice expressing YFP under the control of the CD11c promoter resulting in YFP expression in all CD11c+ cells (24) were used as recipients. hCD2-DsRed mice (gifted by D Kioussis and A Patel, National Institute for Medical Research, London) were crossed with OVA-specific OT-II TCR Tg mice and were used as the T cell donors. Animals were maintained under standard animal house conditions at the University of Glasgow and procedures were performed according to UK Home Office regulations.

### Small molecule library (AZ)

39 compounds were selected from AstraZeneca in-house compound collection. They were all known kinase inhibitors, both from the literature and from in-house AstraZeneca projects and were selected to have varying selectivity profiles towards a diverse range of different kinases in a kinase selectivity panel from The Dundee University.

### Dendritic cell culture

Bone marrow-derived dendritic cells (BMDC) were cultured from C57BL/6 mice as previously described with minor modifications (25). Briefly, femurs and tibia of mice were flushed with complete medium (RPMI-1640 and 10% heat inactivated fetal calf serum (FCS) (Gibco). Single cell suspensions were incubated with 5 mL of red blood cell (RBC) lysis buffer (eBioscience) for 5 min at RT. 1.5×10^6^ cells were seeded in 6-well tissue culture plates (Corning) in 3 mL complete medium supplemented with 20 ng/mL murine recombinant GM-CSF (PeproTech). Cells were incubated in a 37°C and 5% CO_2_ incubator. On day 3 and day 6 complete medium was added or replaced, respectively, with fresh pre-warmed medium supplemented with GM-CSF. On day 7, BMDCs were collected and used for further experiments.

### CD4^+^ T cell enrichment

Spleens and lymph nodes were isolated from OTII TCR Tg mice and single cell suspensions prepared. Spleen cell suspensions were incubated with RBC lysis buffer (eBiosciences) for 5 min at RT. CD4^+^ T cells were obtained by negative selection according to manufacturer’s protocol (130-104-454, Miltenyi Biotec).

### T cell interaction with DC using the IN Cell Analyzer 2000

DCs and CD4^+^ T cells were resuspended at a density of 10^7^ cells/ml in RPMI-1640 medium with 1% FCS. DCs were labelled with Cell Tracker™ Red CMTPX (ThermoFisher) and T cells with vibrant CFDA SE (CFSE) cell tracer kit (ThermoFisher) at 7.5 μM, for 10 min in a 37°C incubator. After incubation, cells were washed by centrifugation at 300 x g for 5 min at 4°C three times in complete media. DCs and CD4^+^ T cells were co-cultured in 384-well μ-clear tissue culture treated microplates (Greiner) in complete medium: RPMI-1640 medium supplemented with 10% heat inactivated FCS and final concentrations of 100 U/mL penicillin and 100 μg/mL steptomycin and 2mM L-Glutamine (Gibco). Cells were incubated with OVA peptide (323-339) (pOVA) (Sigma-Aldrich), whole OVA protein (Worthingtons), Concanavalin A (ConA) at 2.5 μg/mL, (Sigma-Aldrich) in the presence of LPS (Sigma-Aldrich), Y-3P antibody (XCell), protein kinase inhibitors (AstraZeneca) or DMSO as a control for 18h at 37°C and 5% CO_2_ as indicated to assess changes in percentage of T cell overlap with DC as a measure for cell interaction. Cell numbers and the concentration of the stimulants and treatments used are as indicated in each experiment.

### Image acquisition using IN Cell Analyzer 2000 (GE Healthcare)

IN Cell Analyzer 2000 is a lamp-based high content analysis system equipped with a large chip CCD camera (Photometrics CoolSNAP K4). The Nikon 10x magnifying objective with flat field and apochromatic corrections (plan Apo), chromatic aberration-free infinity (CF160) and 0.45 numerical aperture (NA) was used. Cells were imaged at 37°C and 2D images of CFSE-labelled CD4^+^ T cells were acquired in the FITC channel using 490/20x excitation, 525/36m emission. CMTPX-labelled DCs in the TexasRed (TR) channel were acquired using 555/25 nm excitation and 605/52 nm emission. One image covering ~25% of the well was acquired. Images acquired by the IN Cell Analyzer 2000 were analysed using the Developer Toolbox software V1.9.2 (GE Healthcare). Data are shown as representative images or the percentage overlap of OTII T cells with DCs.

### In vitro assessment of T cell activation and proliferation

Day 7 DCs and CFSE labelled CD4^+^ T Cells (prepared as above) were co-cultured at a ratio of 1:1 (100,000 of each) in 96 well plates. Cells were stimulated with pOVA at 1 μg/mL or ConA at 2.5 μg/mL and treated with AS703569 solubilised in DMSO at a final concentration of 0.3 μM or DMSO at a final concentration of 0.1%. At 24 and 72 hours cells were harvested and stained with anti-CD4 eFluor450, anti-CD62L-APCeFluor780, anti-CD25-APC and anti-CD69 PERCPCy5.5 (all eBiosciences). Data was acquired using a MACSquant Analyser (MiltenyiBiotec). Flow cytometry data was analysed using FlowJo10 software (TreeStar)

### Multiphoton imaging

Multiphoton imaging was performed with a Zeiss LSM7 MP system equipped with both a 10x/0.3 NA air and a 20x/1.0NA water immersion objective lens (Zeiss) and a tuneable titanium/sapphire solid state 2-photon excitation source (Chamelon Ultra II; Coherent Laser Group). For *in vivo* imaging, animals were anesthetised with 3% isoflurane in 1.5 L/min oxygen, anaesthesia was maintained with the isoflurane between 1.5-2 %, and oxygen at 1.5 L/min. The leg fixed in place using surgical veterinary glue (Vetbond; 3M) and the popliteal LN was surgically exposed. The hind quarters of the mouse was submerged in PBS warmed and maintained at 35-37°C throughout the experiment (26). Videos were acquired for 15-30 minutes at an X-Y pixel resolution 512 × 512 with 2 μm increments in Z. Images were processed and interaction analysed using Volocity 5.5 (Perkin Elmer) after correction for tissue drift.

### Statistics

Statistical significance was determined using the appropriate test as indicated in the text. A p-value of less than or equal to 0.05 was considered significant. IC50’s were calculated from the dose response using a nonlinear variable slope model with a least squares fit. Data was analysed using GraphPad Prism7.

## ACKNOWLEDGMENTS

This work was in part funded by BTCure grant number,

The authors acknowledge the assistance of Scottish Bioscreening Facility and AstraZeneca Compound Management. The authors acknowledge the assistance of the Institute of Infection, Immunity and Inflammation Flow Cytometry Facility at the University of Glasgow.

---

**Supplementary Figure 1.**
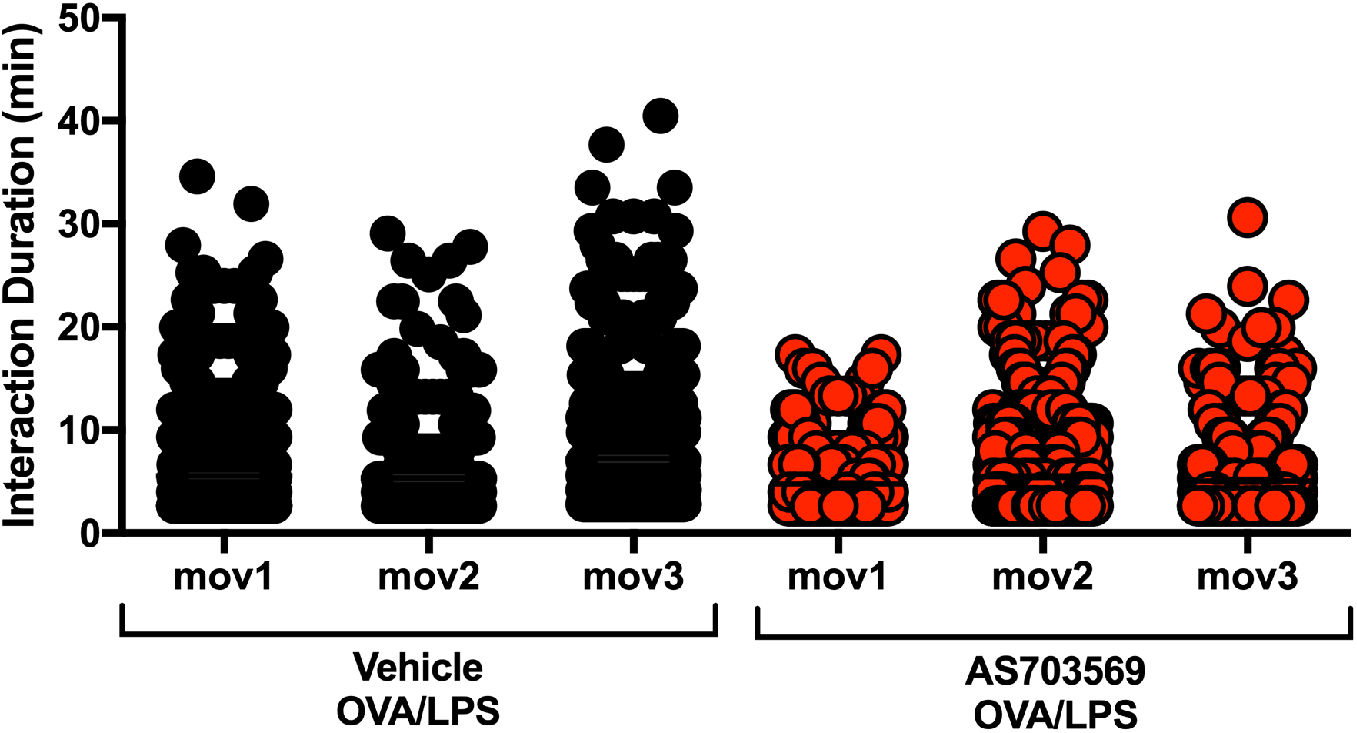
The effect of treatment with AS703569 on T/DC interaction in vivo is short lived. The effect of *in vivo* treatment on T/DC interaction was measured in the lymph nodes of CD11c-YFP mice following the adoptive transfer of DSRed OTII CD4^+^ T cells and subsequent immunisation with OVA/LPS and treatment with 5 μg of AS703579 or vehicle control in the footpad. Three consecutive movies were acquired the Vehicle and AS703569 treated mice (n=1) and the duration of T/DC interaction analysed, each point represents an individual cell. Each movie was approximately 30 minutes duration and the interval between the start of each movie was between 40 and 60 minutes.

*Supplementary Movie 1* Representative movie of popliteal lymph node in OVA/LPS challenged vehicle treated mice, DCs are shown in green and T cells in red, scale bar = 100 μm.

*Supplementary Movie 2* Representative movie of popliteal lymph node in OVA/LPS challenged AS703569 treated mice, DCs are shown in green and T cells in red, scale bar = 100 μm.

## REFERENCES

1. Benson RA, et al. (2015) Antigen presentation kinetics control T cell/dendritic cell interactions and follicular helper T cell generation in vivo. Elife 4:1–16.

2. Zinselmeyer BH, et al. (2005) In situ characterization of CD4+ T cell behavior in mucosal and systemic lymphoid tissues during the induction of oral priming and tolerance. J Exp Med 201(11):1815–23.

3. Mempel TR, Henrickson SE, Von Andrian UH (2004) T-cell priming by dendritic cells in lymph nodes occurs in three distinct phases. Nature 427(6970):154–9.

4. Stephen LA, et al. (2018) The Ciliary Machinery Is Repurposed for T Cell Immune Synapse Trafficking of LCK. Dev Cell 47(1):122–132.e4.

5. Yokosuka T, et al. (2008) Spatiotemporal Regulation of T Cell Costimulation by TCR-CD28 Microclusters and Protein Kinase C θ Translocation. Immunity 29(4):589–601.

6. Brownlie RJ, Zamoyska R (2013) T cell receptor signalling networks: Branched, diversified and bounded. Nat Rev Immunol 13(4):257–269.

7. Serrador JM, et al. (2004) HDAC6 deacetylase activity links the tubulin cytoskeleton with immune synapse organization. Immunity 20(4):417–428.

8. Comrie WA, Li S, Boyle S, Burkhardt JK (2015) The dendritic cell cytoskeleton promotes T cell adhesion and activation by constraining ICAM-1 mobility. J Cell Biol 208(4). doi:10.1083/jcb.201406120.

9. Li L, et al. (2017) Ionic CD3-Lck interaction regulates the initiation of T-cell receptor signaling. Proc Natl Acad Sci U S A 114(29):E5891–E5899.

10. Palacios EH, Weiss A (2004) Function of the Src-family kinases, Lck and Fyn, in T-cell development and activation. Oncogene 23(48 REV. ISS. 7):7990–8000.

11. Bennett SRM, Carbone FR, Karamalis F, Miller JFAP, Heath WR (1997) Induction of a CD8+ Cytotoxic T Lymphocyte Response by Cross-priming Requires Cognate CD4+ T Cell Help. J Exp Med 186(1):65–70.

12. Schoenberger SP, Toes REM, Voort EIH van der, Offringa R, Melief CJM (1998) T-cell help forcytotoxic Tlymphocytes ismediated byCD40–D40L interactions. Nature 255(5505):243–244.

13. Ridge JP, Di Rosa F, Matzinger P (1998) A conditioned dendritic cell can be a temporal bridge between a CD4+ T-helper and a T-killer cell. Nature 393(6684):474–8.

14. Platt AM, et al. (2010) Abatacept limits breach of self-tolerance in a murine model of arthritis via effects on the generation of T follicular helper cells. J Immunol 185(3):1558–67.

15. Emery P (2012) Optimizing outcomes in patients with rheumatoid arthritis and an inadequate response to anti-TNF treatment. Rheumatology (Oxford) 51 Suppl 5:v22–30.

16. Cagnotto G, et al. (2020) Abatacept in rheumatoid arthritis: Survival on drug, clinical outcomes, and their predictors - Data from a large national quality register. Arthritis Res Ther 22(1):1–11.

17. Dinis VG, et al. (2020) Abatacept induced long-term non-progressive reduction in gamma-globulins and autoantibodies: dissociation from disease activity control. Clin Rheumatol (455). doi:10.1007/s10067-020-04932-9.

18. Sharma P, Allison JP (2015) The future of immune checkpoint therapy. Science (80-) 348(6230):56–61.

19. Andersson J, Stefanova I, Stephens GL, Shevach EM (2007) CD4+CD25+ regulatory T cells are activated in vivo by recognition of self. Int Immunol 19(4):557–566.

20. McLaughlin J, et al. (2010) Preclinical characterization of Aurora kinase inhibitor R763/AS703569 identified through an image-based phenotypic screen. J Cancer Res Clin Oncol 136(1):99–113.

21. Blas-Rus N, et al. (2016) Aurora A drives early signalling and vesicle dynamics during T-cell activation. Nat Commun 7:1–16.

22. Keren Z, Berke G (1984) Selective binding of concanavalin A to target cell major histocompatibility antigens is required to induce nonspecific conjugation and lysis by cytolytic T lymphocytes in lectin-dependent cytotoxicity. Cell Immunol 89(2):458–477.

23. Barnden MJ, Allison J, Heath WR, Carbone FR (1998) Defective TCR expression in transgenic mice constructed using cDNA-based α- and β-chain genes under the control of heterologous regulatory elements. Immunol Cell Biol 76(1):34–40.

24. Lindquist RL, et al. (2004) Visualizing dendritic cell networks in vivo. Nat Immunol 5(12):1243–1250.

25. Inaba K, et al. (1992) Generation of large numbers of dendritic cells from mouse bone marrow cultures supplemented with granulocyte/macrophage colony-stimulating factor. J Exp Med 176(6):1693–702.

26. Gibson VB, et al. (2012) A novel method to allow noninvasive, longitudinal imaging of the murine immune system in vivo. Blood 119(11):2545–51.

